# A 96-Well Polyacrylamide Gel for Electrophoresis and Western Blotting

**DOI:** 10.1101/2024.05.10.593650

**Authors:** Marc R. Birtwistle, Jonah R. Huggins, Cameron O. Zadeh, Deepraj Sarmah, Sujata Srikanth, B. Kelly Jones, Lauren N. Cascio, Delphine Dean

## Abstract

Western blotting is a stalwart technique for analyzing specific proteins and/or their post-translational modifications. However, it remains challenging to accommodate more than ~10 samples per experiment without substantial departure from trusted, established protocols involving accessible instrumentation. Here, we describe a 96-sample western blot that conforms to standard 96-well plate dimensional constraints and has little operational deviation from standard western blotting. The main differences are that (i) submerged polyacrylamide gel electrophoresis is operated horizontally (similar to agarose gels) as opposed to vertically, and (ii) a 6 mm thick gel is used, with 2 mm most relevant for membrane transfer (vs ~1 mm typical). Results demonstrate both wet and semi-dry transfer are compatible with this gel thickness. The major tradeoff is reduced molecular weight resolution, due primarily to less available migration distance per sample. We demonstrate proof-of-principle using gels loaded with molecular weight ladder, recombinant protein, and cell lysates. We expect the 96-well western blot will increase reproducibility, efficiency (cost and time ~8-fold), and capacity for biological characterization relative to established western blots.

## Introduction

Western blotting, despite an old-fashioned reputation, remains one of the most (if not the most) widely used protein analysis techniques in biomedical and biological sciences across academia and industry^1,2^. The technique generally uses polyacrylamide slab gel electrophoresis to separate proteins in cell or tissue lysates by molecular weight, and then transfers those separated proteins onto a membrane for subsequent immunodetection. It has also been at the center of multiple recent scientific misconduct controversies^3,4^. While a common first reaction is to suspect some amount of blame is attributable to the technique itself, another and perhaps even likely explanation is that the very nature of western blotting facilitates the identification of misconduct. The reason for that alternative interpretation is the same reason why western blotting is often considered a trusted gold-standard and confirmatory assay: it separates proteins by molecular weight to increase confidence the signal obtained is from the intended target, and then produces a characteristic image of target protein bands that is as yet difficult to fabricate.

Some of the old-fashioned reputation is well-deserved, with western blotting being typically practiced nearly identically to that of its first description over 40 years ago^5–7^. One main limitation of western blotting is being typically restricted to analysis of ~10 samples per experiment. While many intended western blot experiments are not hindered by such sample throughput limitations, others may be. Such demand has led to development of myriad western-like techniques with higher-throughput. The Dodeca cell simply runs up to 12 traditional gels in parallel^8^. Capillary electrophoresis instruments^9^ enable automated analysis of up to 96 samples with groups of 12 simultaneously, but are more expensive compared to western blotting, generate a pseudo blot image derived from chromatography peaks, and may not be as flexible as the western blot with respect to input lysate and buffer types. The microwestern blot enables 96 complete western blots to be performed simultaneously using piezoelectric pipetting to spot small, nL amounts of lysate onto a typical-sized gel^10–12^. Obstacles to microwestern adoption include the piezoelectric pipetting apparatus (expensive, difficult to use, sample loss in tubing, restricted to non-standard lysis buffers, flat gel with no wells), or the use of semi-dry electrophoresis versus the typical immersed tank (expense and difficulty of use). The mesowestern blot^13^, in part developed by us, has the same scale as the microwestern but eliminates the need for piezoelectric pipetting. Yet, it still uses semi-dry electrophoresis, and requires small (~1 μL) sample sizes, which although can be a benefit, is also a detriment if a low abundance target is of interest. Overall, a higher-throughput western blot technique that keeps the familiarity and low operation cost of traditional western blotting could find use and impact.

Here, building on the mesowestern blot, we describe a novel 96-well western blotting technique that has minimal deviation from established western blotting protocols. A key novel component is a slab polyacrylamide gel casted with 96 wells in the geometric format of a 96 well plate, facilitating consistency and familiarity with this widely used standard. The gel is designed to be loaded and subjected to electrophoresis in a submerged horizontal format, similar to agarose gel electrophoresis—this is the primary difference between the described technique and traditional western blotting. To enable analysis of typical lysate volumes, which could be important for low abundance analytes, the gel is 6 mm thick, which gives 56 μL wells, easily capable of holding the typical 20-30 μL of lysate. To simultaneously minimize potential problems caused by gel thickness upon transfer of proteins from the gel to a membrane, the gel is only 2 mm thick from the bottom of the well to the bottom of the gel. Our results demonstrate this gel thickness is not an impediment to successful western blotting with wet or semi-dry transfer. The main tradeoff between the 96-well western blot and traditional western blotting is molecular weight resolution. The primary factor in this tradeoff is simply the reduced distance available in the gel for each sample to travel, but other factors include yet-to-be-developed stacking gel portions or polyacrylamide gradients. To demonstrate proof-of-principle we perform western blot experiments with molecular weight ladder, recombinant protein, and cell lysates. This is the first demonstration of the technique, and we expect substantial room for optimization in subsequent studies.

## Results and Discussion

### Setup and Approach

The microwestern^10–12^ and mesowestern^13^ blot technologies use a horizontal polyacrylamide gel and electrophoresis arrangement to achieve higher sample throughput, and we built on the same approach here. However, these previous techniques required small sample sizes, which are undesirable for medium-to-low abundance targets, and semi-dry electrophoresis, which requires somewhat expensive equipment that is not typically used in western blotting. To address these issues, we designed a casting mold that generates gels with 96, ~56 μL wells (**Fig. 1A-B**). While this is lower throughput than the microwestern and mesowestern (~300 samples), it has the benefit of still being ~10-fold greater throughput than typical 10-12 well vertical arrangements but holding similar sample volume. It was also designed to have geometry matching the widely used 96-well plate standard. We expect this could be useful in future efforts with multi-channel micropipettes or even automation. Lastly, we expect this developed format may be broadly useful to include more replication on the same blot, facilitate direct quantitative comparisons between many samples, include internal recombinant protein calibration curves in every experiment, explore more experimental conditions, or to probe the same set of samples with different antibodies more efficiently by cutting the membrane into multiple sections^14,15^.

**Figure 1.**
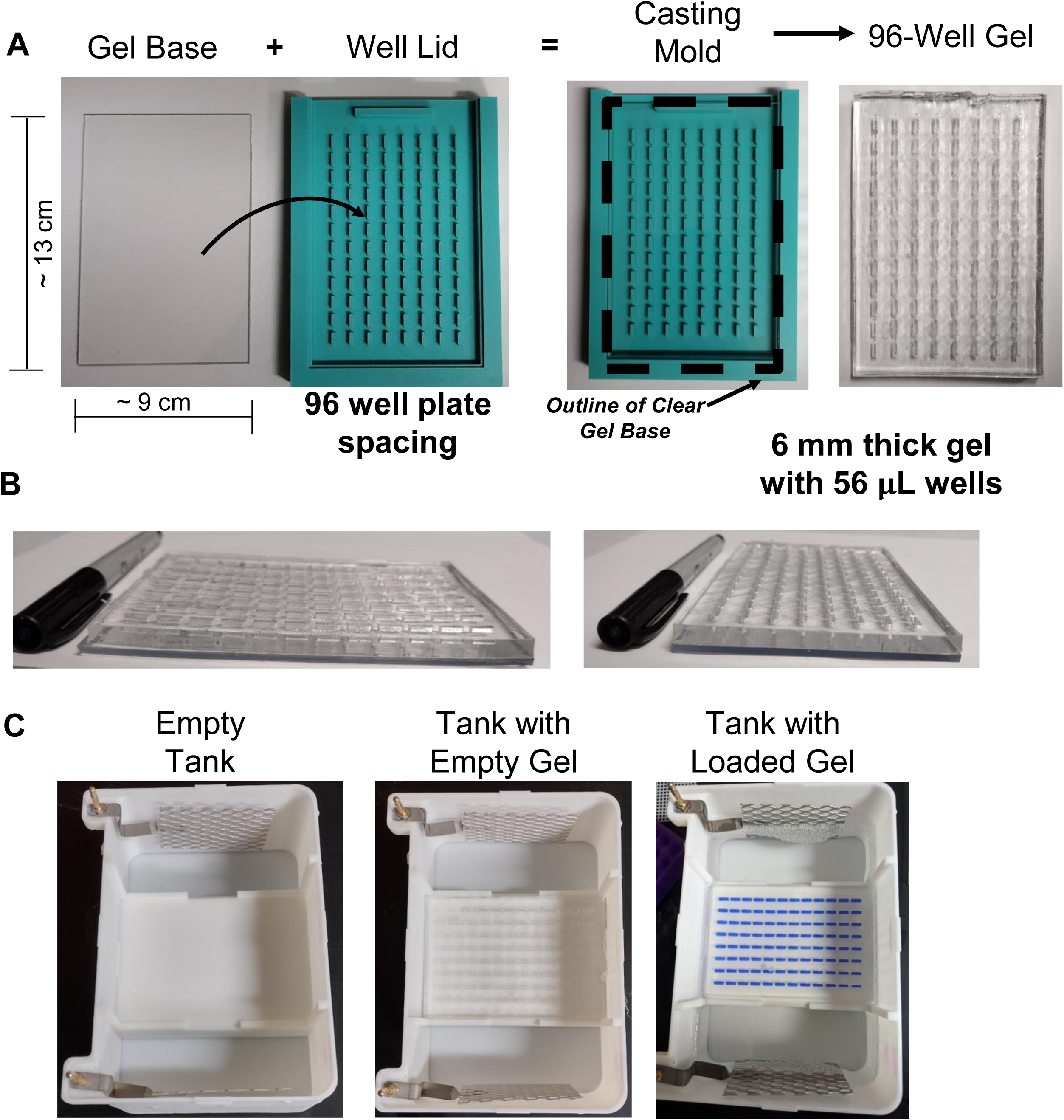
New Apparati Enabling the 96-Well Western Blot. **A.** Gel casting mold. The ABS “Well Lid” imprints the wells into the eventual gel, and the polycarbonate “Gel Base” serves as the gel support after casting. The two pieces fit together to enable casting (after clamping and sealing). **B**. Long edge (left) and short edge (right) of gel, side views. The thin sharpie pen is for perspective and reference. **C.** Horizontal electrophoresis tank with no gel (left), empty gel (center), and loaded gel (right). The gel base and gel are placed onto the center tray in the tank. The tank is then filled with running buffer, the gel loaded with samples, and finally subjected to electrophoresis. After that, transfer is standard.

The biggest tradeoff for using this 96-well gel versus traditional vertical gels is reduced molecular weight resolution. In some, and perhaps even many situations, the full degree of molecular weight resolution afforded by traditional blotting may not be needed, although we expect determination of sufficiency to be highly context dependent. There are three main factors driving the tradeoff. First, an almost certainly irreconcilable factor is that each sample has roughly 1/8 the distance for separation as compared to a traditional vertical gel, simply due to stacked rows of wells. Second, the current casting design precludes addition of a small low polyacrylamide % (and pH) “stacking” gel which is generally thought to improve molecular weight resolution^16–18^. Third, the current casting design precludes generation of an acrylamide gradient for each well, which when included makes migration distances for different size proteins more uniform^19^. We expect that further innovation could address the second and third factors. However, to simply establish proof-of-principle in this work, we focus only on 10% acrylamide gels. Acrylamide % is simple to change if different size proteins need to be resolved.

The gel is designed for a submerged horizontal electrophoresis tank (**Fig. 1C**), similar to agarose gels for nucleic acids^20^. Not only does such a gel enable loading of typical sample sizes (we used between 5 μL and 20 μL in this study), it also greatly reduces the possibility of sample washout, since the gel is loaded with dense, sinking sample (due to glycerol content) while already submerged with electrophoresis running buffer. In our experience so far, we did not experience any issues with sample washout or difficulty reproducibly loading samples in these gels. While we do have a specific tank design that is shown to work here, it is not unique, and we expect other designs that hold the gel would also work.

### Molecular Weight Ladder Experiments

As a first test of this system, we loaded 10% acrylamide gels with two different molecular weight ladder standards—infrared fluorescent and colorimetric, and then subjected them to transfer, both wet and semi-dry (**Fig. 2**; **Fig. S1**). Transfer did not seem to be impeded despite the gel being thicker than normal. As expected, the ladder components had reduced resolution compared to that normally observed in such blotting, primarily due to the reduced available migration distance. However, clear bands were observed in ranges between ~25 kDa and ~100 kDa for this 10% gel, as expected. We noted that bands greater than 150 kDa in the colorimetric ladder experiment remained in the gel, and/or possibly did not appreciably enter the gel from the well (gel images were not available). Using 6% acrylamide gels enabled resolution of MW bands between 100 and 250 kDa (**Fig. 2C**). On the lower end of MW, in 10% acrylamide gels, proteins <~25 kDa yielded diffuse bands. Analogously, 20% acrylamide gels enabled resolution of MW bands between 10 and 37 kDa (**Fig. 2D**). We conclude that 10%, 96-well polyacrylamide gels can effectively resolve proteins in the ~25 – 100 kDa range and proteins from these gels can be robustly transferred to membranes. Moreover, gels with lower or higher acrylamide % modulate the resolvable MW range in expected ways. Taken together, these results demonstrate that 96-well polyacrylamide gels can resolve proteins between 10 and 250 kDa.

**Figure 2.**
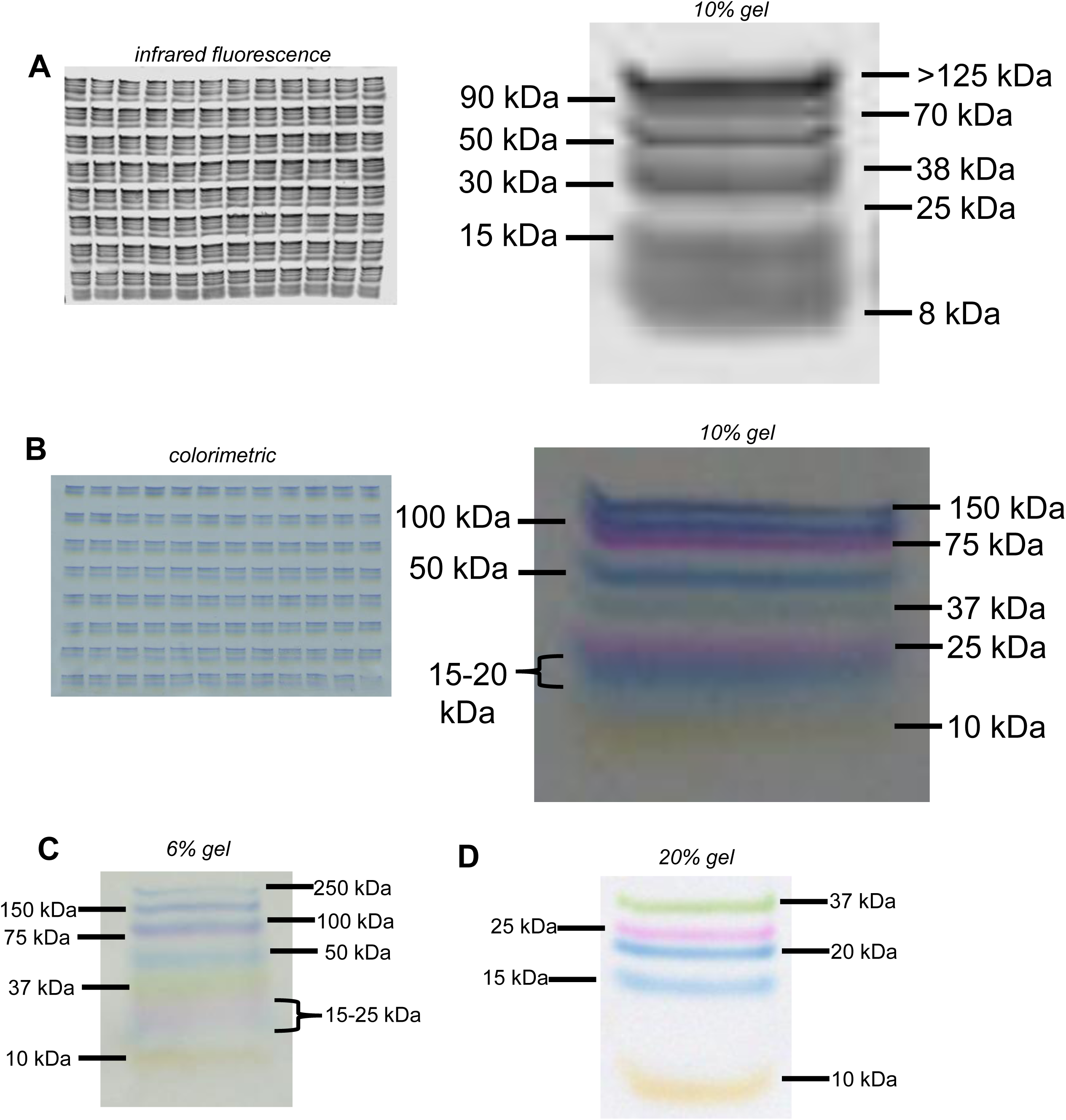
96-Well Blot with Molecular Weight Ladder. **A.** Infrared fluorescence molecular weight ladder experiment. A 96-well 10% Tris-HCl gel was loaded with 4.5 *μ*L of molecular weight ladder, subjected to electrophoresis, transferred to nitrocellulose membrane, and imaged. Left: Full membrane scan. Right: Example lane. Different molecular weight bands from the ladder are indicated. **B.** Colorimetric molecular weight ladder experiment. A 96-well 10% Tris-HCl gel was loaded with 5 *μ*L of molecular weight ladder, subjected to electrophoresis, transferred to PVDF membrane, and imaged. Left: Full membrane scan. Right: Example lane. Different molecular weight bands from the ladder are indicated. For all panels, full membrane scans for replicate experiments and those involving semi-dry transfer are presented in Fig. S1. **C-D.** Example lane from a 6% gel (C) or 20% gel (D). Experiment was done as in (B) but with nitrocellulose membrane.

### Recombinant Protein Experiments

Next, we wondered how reproducible the system might be with regards to band signal intensities. To test this, we loaded 20 μL (25 ng) of α-Tubulin into each well in the middle 10 columns of a 10% gel (outside 2 reserved for molecular weight ladder), and performed western blotting using chemiluminescent detection (**Fig. 3A**, **Fig. S2**). Overall, molecular weight resolution was reasonably uniform across the entire membrane, with single bands observable at the expected molecular weight (~50 kDa). Visual band density appeared overall uniform with expected variation. In replicate experiments (**Fig. S2B**), transferred using a faster protocol (see Methods), we noted potentially lower transfer efficiency on some membrane edges. Given this many samples run simultaneously, it is not unexpected to have a few systematic deviations, likely due to transfer. This highlights the importance of including an internal (loading) control for each lane, which is most commonly a “housekeeping” protein such as α-Tubulin, but can include a variety of total protein normalization methods^21^.

**Figure 3.**
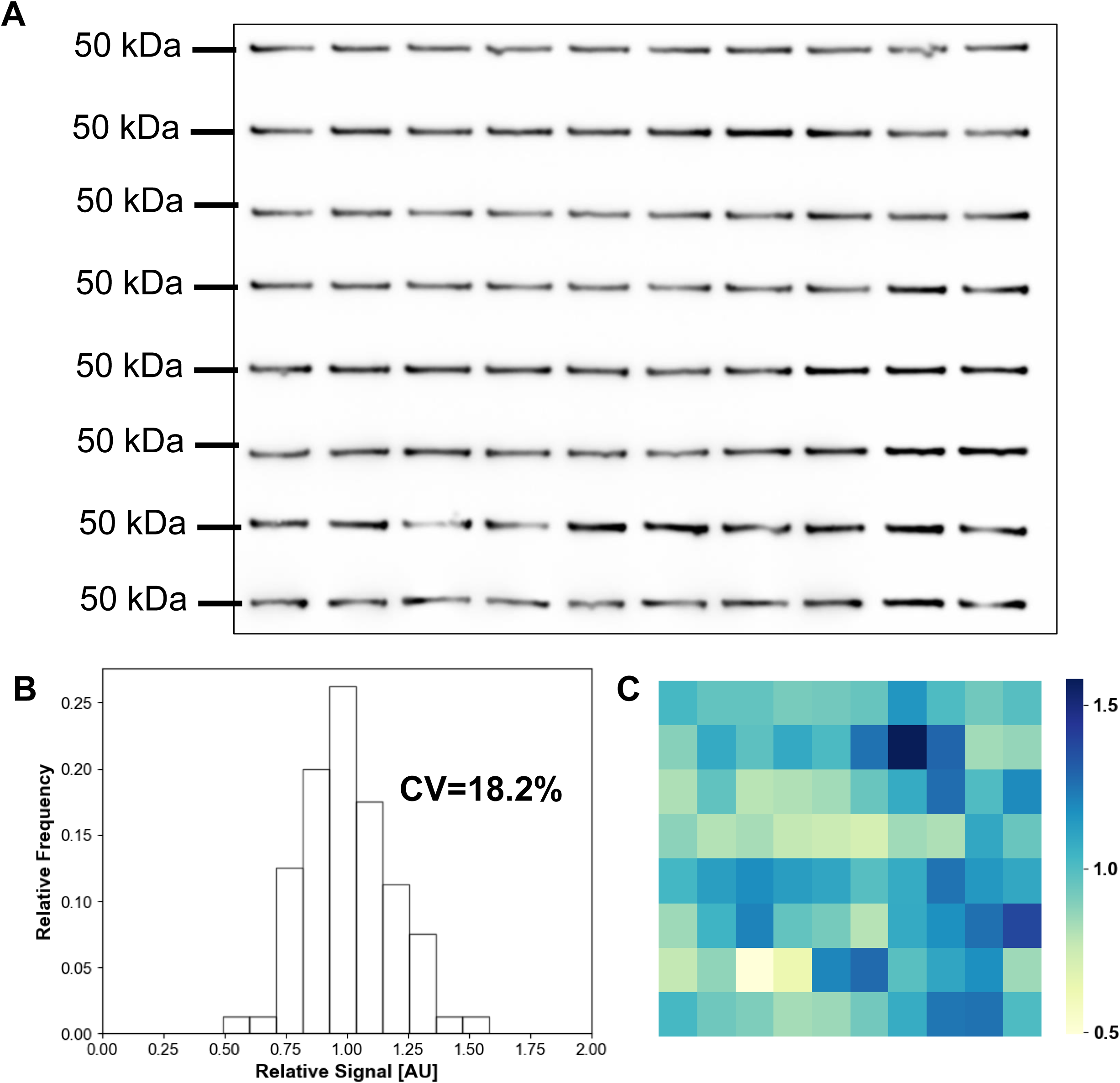
96-Well Western Blot Loaded with Recombinant Protein. **A.** Full membrane scan. A 96-well 10% Tris-HCl gel was loaded with 5 *μ*L of molecular weight ladder in each well of the first and last columns, and 20 *μ*L / 25 ng of recombinant *α*-Tubulin (~50 kDa) in the other wells. We then performed electrophoresis, transfer (30V, 16 hr, 4°C, nitrocellulose), and antibody incubations, and finally imaged. Replicate experiment full scans are in Fig. S2 (along with combined ladder scans for the presented result). **B.** The bands from A were subjected to densitometry and the relative intensities analyzed for variation as a histogram. **C**. Densitometry results of band quantification from image in A. Each band corresponds to a square in the heatmap.

Quantification of the band intensities showed a visually symmetric distribution (**Fig. 3B**) with some spatially correlated variation (**Fig. 3C**). This is not unexpected since a major source of quantitative variation in western blotting is typically spatial due to transfer heterogeneity. The coefficient of variation (CV) was 18.2%. In a similar experiment with cell lysate instead of recombinant protein, the CV was comparable at 16.1% (**Fig. S3**). This would allow for reliable detection (~90% confidence, ~80% power) of ~1.5 fold changes with triplicate measurements. Previous western blotting work reports CV from 15% to 70% (or more)^13,22–25^, so this value is typical or even low for western blots. Importantly, the CV reported for this particular 96-well western is artificially high since there is by definition no loading control, which is generally known to improve CV^21^ by about a factor of 1.25^13^, down to ~12-20%^24,25^. We performed dual-color infrared fluorescence imaging with GAPDH and α−Tubulin (**Fig. S4**), and found loading control normalization improves CV by a factor of 1.17, consistent with prior work. Conveniently, one potential use of these 96-well blots is to include replication in the gel loading design to improve statistical performance. We conclude that the 96-well gel enables transfer of protein to membranes for immunodetection with reproducibility at least equal to that of traditional western blots.

### Recombinant Protein and Cell Lysate Serial Dilution Experiments

Next, we wanted to test the 96-well gel using 2-fold serial dilutions of both recombinant protein and cell lysate, again probing for α-Tubulin (**Fig. 4**). This is expected to be a potential application of the 96-well western blot for many researchers, to do multiple optimizations at scale with a single gel. To do this, we loaded a 10% gel with varying concentrations of either recombinant α-Tubulin or HEK293 cell lysate, and performed western blotting using chemiluminescent detection. As above, we observed reasonably uniform molecular weight separation across the gel with identifiable bands at the expected molecular weight (~50 kDa). We also observed dose dependent band intensities visually, which matched expectations based on loading patterns. Quantified band intensities revealed somewhat small linear ranges with hyperbolic saturation, typical for chemiluminescent detection^21^. The average CV across replicates (for samples above detection limit) for recombinant α-Tubulin samples was 22.2%, comparable to above, especially given the smaller number of replicates here versus above. Interestingly, the average CV across replicates for cell lysate samples was 5.0%, significantly lower. We also noted the cell lysate bands were less diffuse, which we speculate may be caused by the presence of other proteins. Thus, although much more extensive testing is required, the expected CV for cell lysate samples, the true eventual application for most researchers, may be quite low. We would also note here that as above, there is no housekeeping or total protein control, which would likely lower CV. Lastly, we do not perform extensive analysis of linear range or limit-of-detection, because detection modality and antibody epitope are the primary drivers, and these are routinely changed depending on the intended investigation. Importantly, however, the observed limit-of-detection is essentially equivalent to that observed in an analogous western blot experiment (**Fig. 4D**).

**Figure 4.**
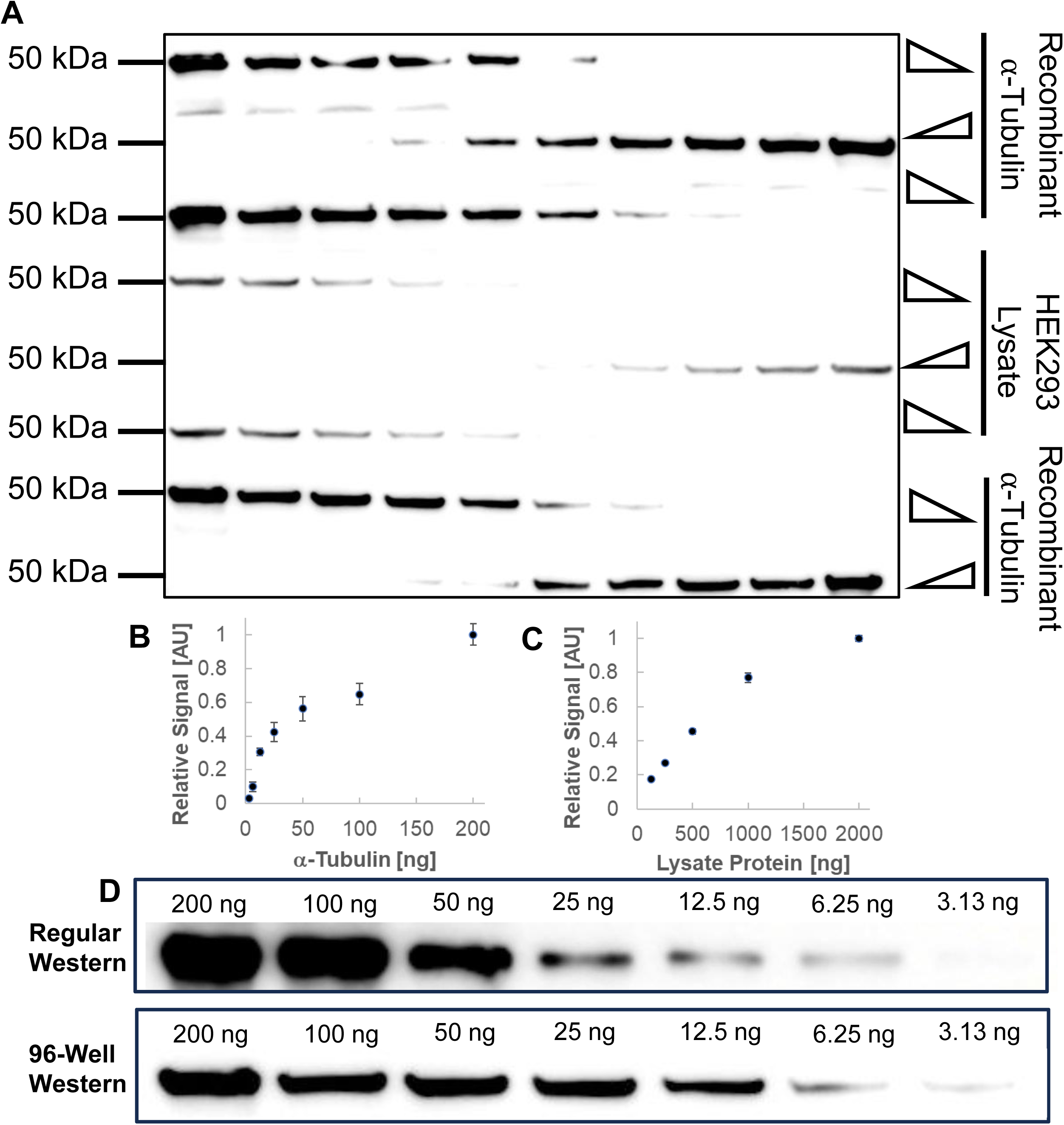
96-Well Western Blot with Lysate and Recombinant Protein Serial Dilutions. **A.** Full membrane scan. A 96-well 10% Tris-HCl gel was loaded with 5 *μ*L of molecular weight ladder in each well of the first and last columns (not shown). 4 *μ*L of either recombinant *α*-Tubulin or HEK293 lysate was loaded in the other wells in 2-fold dilution series as indicated. We then performed electrophoresis, wet transfer (PVDF), antibody incubations, and imaged via chemiluminescence. **B-C.** The bands from A for recombinant protein (B) and lysate (C) were subjected to densitometry and the relative intensities quantified. Error bars denote standard error from replicates on this blot. **D.** Comparison of regular western to 96-well western (7^th^ row above). The same amount and types of recombinant protein (α-tubulin), antibodies, and chemiluminescence substrate were used.

### Conclusions

We present a 96-well western blot that has high similarity to traditional western blotting, with the main difference being submerged horizontal rather than submerged vertical electrophoresis. The tradeoff is molecular weight resolution, caused primarily by simply less distance for proteins to migrate. We demonstrate proof-of-principle with multiple experiments involving molecular weight ladder, recombinant protein, and cell lysate. Quantitative performance is comparable to previously reported traditional western blot results. We expect this technique to be valuable to a broad variety of academic and industrial researchers investigating dose and dynamic responses, biomarkers across multiple conditions or samples, and internal, same-membrane replication for increased quantitative fidelity and reproducibility. Moreover, we expect substantial time and cost savings with this system compared to regular western blot, approximately 85% reduction in consumables cost (gels, membranes, blotting paper, antibodies, substrates) and ~8x efficiency for time savings.

## Methods

### Casting Gels

Additive manufacturing (Raise 3D Pro3) with ABS filament (Polymaker, #PE01010, Amazon) was used to construct the well lid. After printing, the bottom side of the well lid was sprayed with clear Flex Seal (#FSCL20, Lowe’s). Polycarbonate sheets (0.093 in thick, #1PC0081A, Lowe’s) were cut to size (128 mm by 86 mm) with a table saw to construct gel bases. The gel base was placed into the well lid (**Fig. 1**) to create the casting mold. A 6”x6”x1/16” rubber mat (Keeney, #PP25546, Lowe’s) was placed on top of the gel base side of the casting mold, and subsequently two cut-to-size pieces of 5/8” thick plywood were placed on either side of the casting mold. The entire assembly was then placed vertically into a woodworker’s vice (PONY, #26545, Lowe’s) with the loading edge upwards, with only slight pressure. Two additional expandable clamps (Jorgensen, #93366, Lowe’s) were placed and tightened on the loading edge vertically aligned, and then the bottom woodworker’s vice was fully tightened. The order of tightening helps ensure excess bottom pressure does not force the gel base upwards prior to clamping the top.

Unpolymerized 10% Tris-HCl gel solution (250 mL) was prepared by combining 30% bis/acrylamide (83.33 mL, #1610156, BIO-RAD), 1.5M Tris-HCl, pH 8.8 (62.5 mL, recipe below), 10% (g/100 mL) SDS (2.5 mL, recipe below), with water (Primo, Walmart) to 250 mL. Different % gels were made by scaling the amount of bis/acrylamide proportionally. Lower % gels included 5% (v/v) glycerol (#AAA16205AP, ThermoFisher). The 1.5M Tris-HCl solution was made by dissolving 181.5 g of Tris (#140500025, ThermoFisher) in ~500 mL of water, then drop dispensing concentrated HCl (#389310025, ThermoFisher) while monitoring pH with continuous magnetic stirring until pH=8.8, then adding water to 1000 mL. 10% SDS was made by dissolving 10 g of SDS (#28312, ThermoFisher) in 80 mL of water with gentle magnetic stirring, then bringing to 100 mL with water. To prepare for gel casting, 70 mL of unpolymerized gel solution, 350 μL of 10% (g/100 mL) ammonium persulfate (APS, recipe below), and 35 μL of TEMED (#1610801, BIO-RAD) were added to a glass beaker and mixed thoroughly. The 10% APS solution was made by dissolving 1 g of APS (#1610700, BIO-RAD) in 10 mL water and storing at 4°C. Serological pipettes were used to transfer the unpolymerized gel solution with APS and TEMED into the clamped casting assembly slowly. Gels were allowed to polymerize for at least 6 hours but no more than 12 hours. Gels were removed from the casting assembly carefully, a plastic lid placed over the wells, and then placed into a vacuum sealing bag (Wevac, B07TV5RNQL, Amazon) with 5 mL of electrophoresis running buffer (recipe in below section), and sealed (Nesco, VS-12, Amazon). Gels were stored at 4°C and used within a month.

### Horizontal Electrophoresis

Additive manufacturing (Raise 3D Pro3) with Hyper Speed ABS filament (Raise3D) was used to construct the electrophoresis tank. After printing, the bottom and sides of the tank were sprayed with clear Flex Seal (#FSCL20, Lowe’s). Banana plugs (Guangdong Techeng Hardware Electronics, #BP2778-A, Alibaba) and platinized titanium meshes (PLANODE2X3, Amazon) were integrated into the tank prior to use.

A gel cast as above (#G96, Blotting Innovations, LLC) was removed from the vacuum sealed package, and then placed with the gel base into the center tray area of the electrophoresis tank (#T1, Blotting Innovations, LLC) on a level surface. The tank was filled with enough cold (4°C) Tris-HCl running buffer to cover the gel (~600 mL), and then the gel was loaded with sample. Tris-HCl running buffer (1 L) was prepared by dissolving 12.1 g of Tris in ~500 mL water, drop dispensing HCl as above but instead to pH 7.5, adding water to 990 mL, and then 10 mL 10% SDS. A 10X stock solution was also often prepared and used.

Colorimetric molecular weight ladder (Kaleidoscope #1610395, BIO-RAD) was mixed 1:1 (v/v) with clear sample buffer (recipe below) and ~5 μL loaded per well. Clear sample buffer (1 mL) was prepared with 500 μL of 50% (v/v water) glycerol (#AAA16205AP, ThermoFisher), 200 μL 10% SDS (see above), 42 μL 1.5 M Tris-HCl pH 6.8 (as above), 50 μL 2-Mercaptoethanol (#AC125472500, ThermoFisher), and water to 1 mL.

Infrared fluorescent molecular weight ladder (500 μL Chameleon Duo #928-60000, LI-COR) was mixed with 20 μL of 0.1% (g / 100 mL) bromophenol blue loading dye solution (recipe below) and 30 μL of 100% glycerol, and 4.5 μL loaded per well. Bromophenol blue loading dye solution (10 mL) was prepared with 0.01 g bromophenol blue (#AAA1689918, ThermoFisher), 5 mL of 50% glycerol (see above), 0.5 mL of 10% SDS (see above), 2.1 mL of 1.5M Tris-HCl pH 6.8 (see above), and water to 10 mL.

Recombinant α-Tubulin (#ag18034, Proteintech) resuspended in 50% glycerol (see above) to 0.1 mg/mL or HEK293 cell lysate (#CL-07, Protein Biotechnologies) were mixed 1:1 (v/v) with 2X Laemmli sample buffer. The 2X Laemmli sample buffer (45 mL) was prepared with 6.25 mL 1M Tris pH 6.8 (60.55g Tris in 500 mL water, prepared as above except pH to 6.8), 20 mL 10% SDS, 10 mL 100% glycerol, 0.5 mL 1% (g/100 mL) bromophenol blue (0.5 g / 50 mL water), and water to 45 mL. Immediately prior to use, 2-Mercaptoethanol was added to complete the 2X sample buffer at 10% (v/v) (50 μL / 500 μL final). Samples were heated at 95°C for 5 minutes and allowed to cool before loading. If samples were further diluted prior to loading, 2X Laemmli sample buffer was mixed 1:1 (v/v) with water to make 1X Laemmli sample buffer, and then this was used to dilute as appropriate. Gels were loaded with either 5 μL or 20 μL of sample as indicated.

After sample loading, the tank banana plugs were connected to a power source (90W Life Technologies PowerEase, eBay) with banana plug connectors (Longdex, #BA4.0-Y240-RB2, Amazon), and cut-to-size plastic mesh (Sativa, # D05030, Amazon) was placed perpendicular to the tank flanking the gel to prevent foam and bubble build up over the gel during electrophoresis. Newer versions of the T1 tank have this mesh integrated into the lid. Electrophoresis was conducted at 50V for ~45 minutes with monitoring of bromophenol blue dye front, with electrophoresis stopped when this dye front reached the next well. For the 20% acrylamide gel, longer electrophoresis times were needed, which could be mitigated by higher voltages.

### Wet Transfer

Prior to the experiments, wet transfer buffer was prepared by (i) dissolving 30.3 g Tris (#140500025, ThermoFisher) and 144 g glycine (#1610724, BIO-RAD) in 10 L water, and (ii) the day before use, adding 400 mL methanol (MAXTITE, Amazon) to 1600 mL of the above and storing at 4°C to ensure an ample supply of cold buffer. After electrophoresis, the gel with base was transferred into a plastic tray containing cold transfer buffer and allowed to incubate for 10 minutes. A transfer sandwich was constructed using the BIO-RAD Criterion blotter kit (#1704071) and manufacturer’s instructions (**Fig. S5A**). Briefly, cold transfer buffer-soaked blotter paper (#1704085, BIO-RAD) was placed on top of the gel (well side up), and then placed onto a transfer cassette (black side down) with a similarly soaked sponge below (from kit), all immersed in cold transfer buffer. Cut-to-size nitrocellulose (#1620112, BIO-RAD) or PVDF (#1620177, BIO-RAD) membranes were equilibrated in cold transfer buffer (PVDF was pre-wetted in methanol), placed onto the gel (flat side without wells), and carefully rolled (using roller from kit). We obtained better contact between membrane and gel by having the whole assembly submerged in transfer buffer during this stage. Cold transfer buffer-soaked blotter paper was placed on top of the membrane, gently rolled, a soaked sponge placed on top, then the cassette closed. Care was taken to not pinch or damage the gel with cassette protrusions or the sliding clamp near edges. The transfer sandwich was placed into the Criterion blotter in which a magnetic stir bar and ice-pack were placed. The tank was filled with cold transfer buffer, magnetic stirring started, and subjected to either constant 500 mA for 90 minutes at room temperature or 30 V for 16 hours at 4°C.

### Semi-Dry Transfer

Prior to experiments, semi-dry transfer buffer (250 mL) was prepared by dissolving 14.53 g Tris (#140500025, ThermoFisher) and 7.31 g glycine (#1610724, BIO-RAD) in 200 mL water and then adding 50 mL methanol (MAXTITE, Amazon) before use. After electrophoresis, the gel was soaked in transfer buffer for 10 minutes with wells facing up. A transfer sandwich was constructed using 6 transfer buffer-soaked blotter papers (#1704085, BIO-RAD) (**Fig. S4**). First, 3 soaked blotter papers were stacked on top of the gel, still with wells facing up submerged in transfer buffer. This mode of assembly helped to ensure all wells contained transfer buffer with no air, to facilitate efficient and uniform transfer. This half-sandwich was lifted with a spatula, and placed blotter paper side down onto the top of the semi-dry transfer apparatus cassette (cathode), and rolled gently (**Fig. S5B**). Next, a cut-to-size nitrocellulose membrane that was pre-soaked in transfer buffer (#1620112, BIO-RAD) was placed onto the smooth side of the gel, and rolled gently. Next, 3 more transfer buffer-soaked blotter papers were placed on top of the membrane and gently rolled. The bottom of the apparatus (anode) was then attached and the transfer cassette (Powerblotter, Pierce/ThermoFisher, eBay) was sealed firmly. Transfer was conducted at 10V for 50 minutes. In our observations so far, the pressure on the sandwich is a key variable for good and uniform transfer, which is provided by extra blotter papers.

### Membrane Treatment

The membrane was removed from the sandwich and inspected for efficient molecular weight ladder transfer. If only molecular weight ladder was loaded, the membrane was then imaged (see below). If other samples were present, the membrane was immediately placed into a glass dish containing 20 mL of room temperature blocking buffer, and subjected to 100 rpm on an orbital shaker (ONiLAB, #SK-O180-S, Amazon) for 1 hour. Blocking buffer was prepared by adding 5 g non-fat dry milk (Great Value, Wal-Mart) to 100 mL of TBS-T (Tris Buffered Saline with Tween-20, recipe as follows). A 10X TBS stock (1 L) was prepared by dissolving 24 g Tris and 88 g NaCl (#AA12314A3, ThermoFisher) in ~500 mL of water, adjusting pH to 7.6 as above with HCl, and then adding water to 1 L. A 10% (v/v) Tween-20 (#BP337500, ThermoFisher) solution was prepared by adding 10 mL Tween-20 to 90 mL water. A 1X TBS-T solution (1 L) was prepared by mixing 100 mL 10X TBS into 890 mL water, then adding 10 mL of 10% Tween-20. For infrared fluorescence experiments, Tween-20 was not used in the blocking buffer.

After blocking, the membrane was incubated with 20 mL of primary antibody solution at room temperature for 1 hour at 100 rpm. The primary antibody solution was prepared by adding anti-α-Tubulin (#11224-1-AP, Proteintech, Rabbit IgG) 1:5,000 (v/v) to blocking buffer or anti-GAPDH (#60004-1-1g, Proteinetch, Mouse IgG) 1:10,000 (v/v) to blocking buffer. The membrane was then washed 3 times with ~20 mL of TBS-T for 5 minutes at 100 rpm at room temperature. Following washing, the membrane was incubated with 20 mL of secondary antibody solution at room temperature for 1 hour at 100 rpm. Secondary antibody solution was prepared by adding HRP-conjugated Goat Anti-Rabbit IgG (#SA00001-2, Proteintech) 1:10,000 (v/v) in TBS-T or a combination of IR680-conjugated Goat Anti-Mouse IgG (#925-680070, LICOR) and IR800-conjugated Goat Anti-Rabbit IgG (#925-32211, LICOR) 1:10,000 (v/v) in TBS-T. The membrane was then washed 3 times with ~20 mL of TBS-T for 5 minutes at 100 rpm at room temperature. After the final wash, ~ 2 mL of chemiluminescent substrate solution prepared according to manufacturer’s instructions (#PI34577, ThermoFisher) was applied dropwise to the top of the membrane to prepare for imaging.

### Imaging and Analysis

For infrared fluorescent molecular weight ladder experiments, a LI-COR Odyssey was used. The membrane was placed onto the scanner bed and both 700 nm and 800 nm channels were acquired with 169 μm resolution. The resulting image was converted to grayscale for presentation in figures.

For all other experiments, an Azure 300 (#AZ1300-01, Avantor) or 600 was used. The membrane was placed onto the black chemi-tray which was placed onto the top shelf in the unit. Acquisition was done via the Chemi Blot module with automatic exposure time calculation, cumulative image generation, and color marker selected. Any images with saturated pixels were discarded. Images were saved as `.jpg` for figure generation and `.tiff` for quantitative analysis.

For quantitative analysis of chemiluminescent bands, ImageJ was used. A rectanglular region of interest was placed over one row of bands at a time. Under AnalyzeGels, select first lane was chosen, and then plot lanes. On the generated intensity profile plots, vertical lines were drawn manually to separate peaks, and then the wand tool was used by clicking inside each peak to generate the area metric for each band. This process was repeated for each row in a blot image, and arbitrary area units were scaled to be unitless.

### Regular Western

All the above methods were followed for implementation of regular westerns, with the following exceptions. Pre-cast 10% Criterion TGX gels (#5671033, BIO-RAD) were used with a Criterion Electrophoresis Cell (eBay) according to the manufacturer’s instructions. Wet transfer with the Criterion Cell was used (not semi-dry).

## Author Contributions

MRB, COZ, and JRH conceived of the ideas and experiments, and performed and supervised the work. DS performed semi-dry transfer experiments. SS, BKJ, and LNC performed and DD supervised the infrared fluorescence molecular weight ladder experiments. MRB wrote the paper.

## Acknowledgements

We thank NIH / NIGMS (R41GM148112) for funding, Eugene Krenstel at XLerate Health and Tyler Tatum at 3PhaseSC for advice and support, and Kim Schwartz at Advansta for helpful discussions.

---

**Figure S1.**
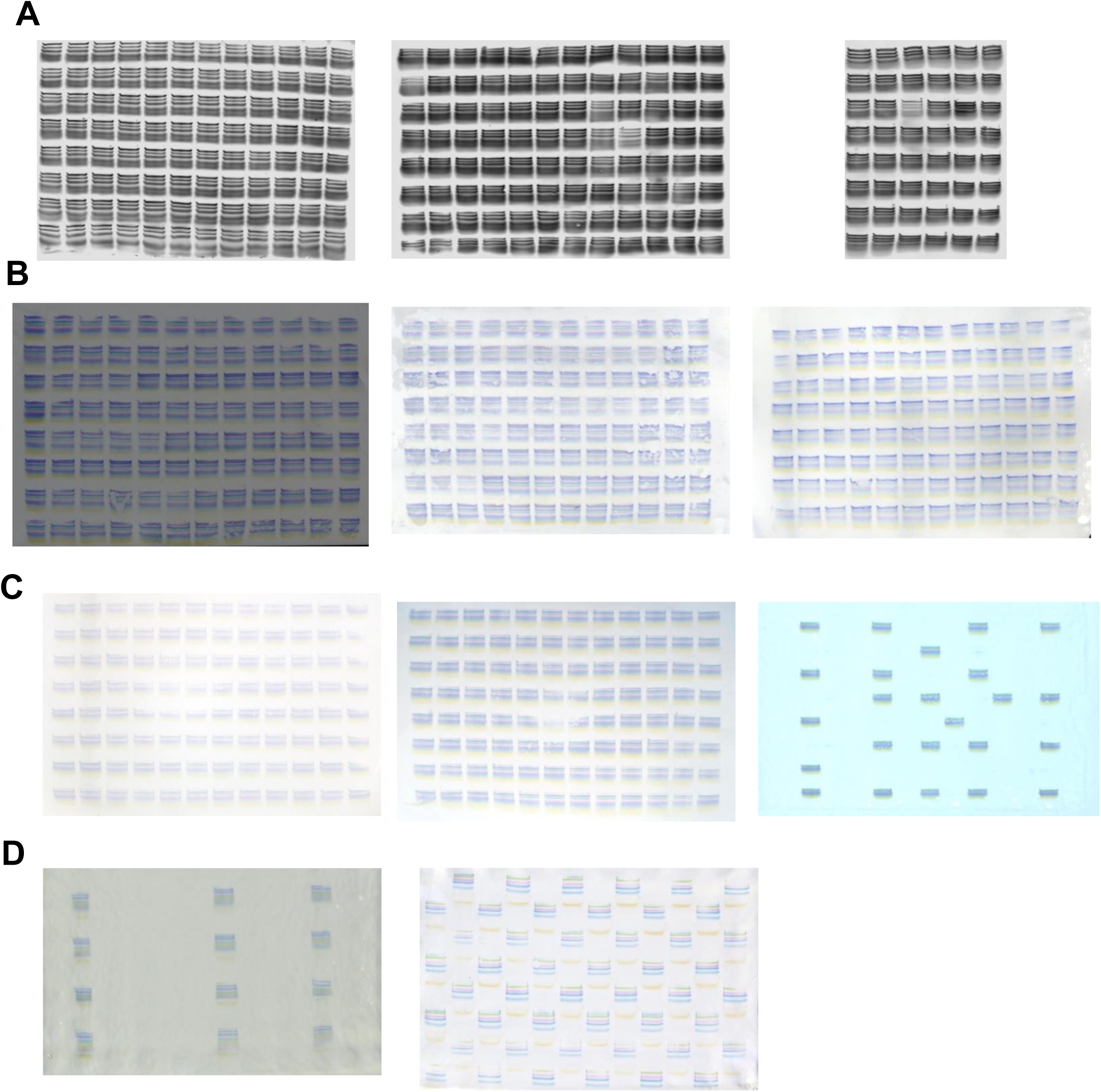
Full Scans of Replicate 96-Well Blots with Molecular Weight Ladder. Experiments were carried out as in Figure 1. **A.** Infrared fluorescence ladder experiments. Only the 700 nm channel is shown for clarity (Chameleon Duo was used which also has bands in the 800 nm channel). The far right result only shows a half gel because wells in the other half of the gel were defective from manufacturing—this did not interfere with transfer of the remaining gel. **B.** Colorimetric ladder experiments. On the far left membrane, the bottom row was close to the top of the transfer cassette and we hypothesize it was not completely submerged in buffer. The far right blot was performed with 30V, 16 hours, 4°C transfer. **C.** Colorimetric ladder experiments using semi-dry transfer. The far right blot had only intermittent loaded ladder. **D**. Full membrane scans for 6% (left) and 20% (right) gels.

**Figure S2.**
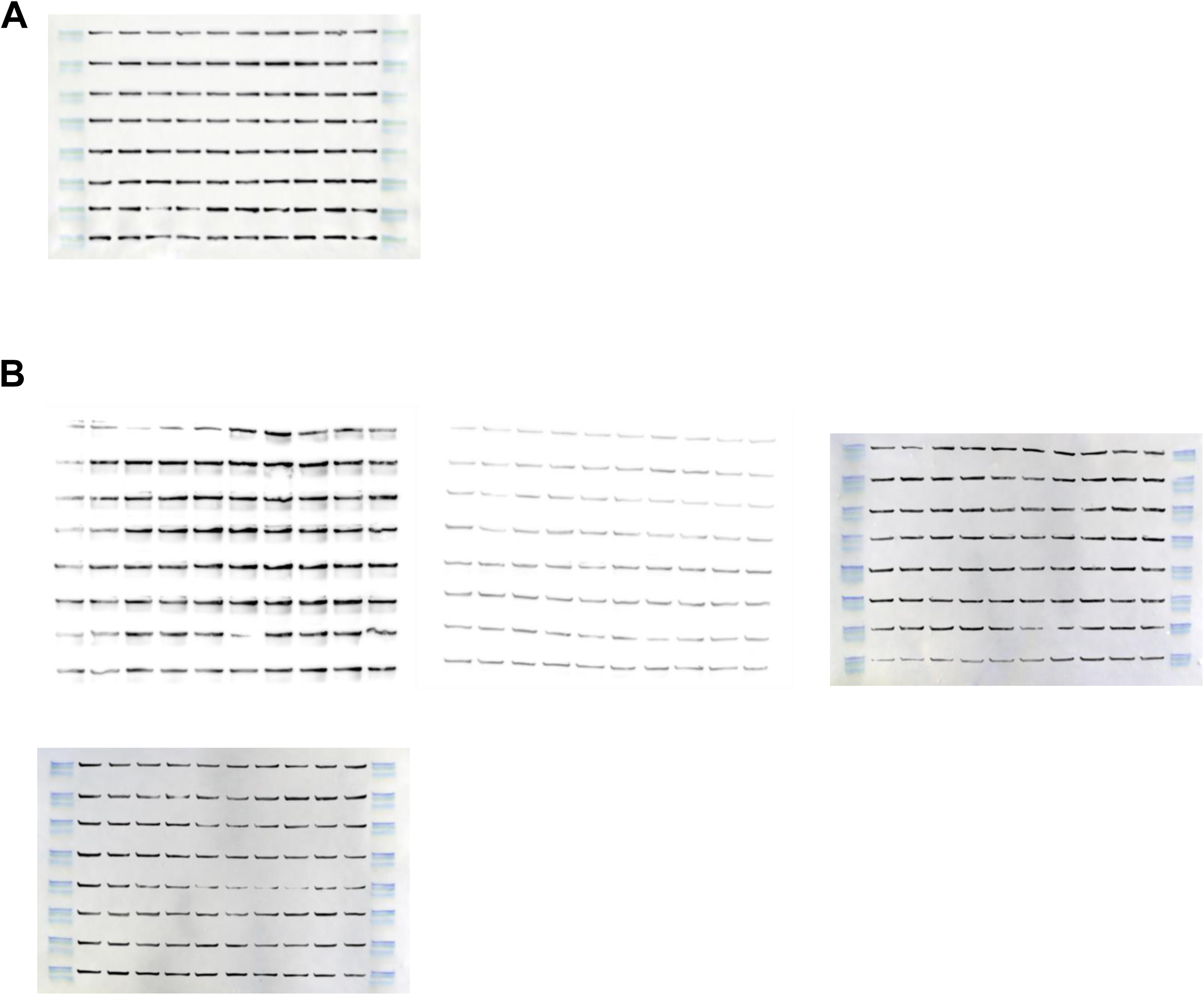
96-Well Gel and Blot Loaded with Recombinant Protein. **A.** Full membrane scan with ladder from the result in Fig. 3. **B.** Independent replicates of the Fig. 3 experiment. The top row were loaded with 5 uL, 25 ng of protein, and transferred at 500 mA for 1.5 hrs. The bottom is the same as panel A, 20 uL, 25 ng of protein, and transferred 16 hrs at 30 V.

**Figure S3.**
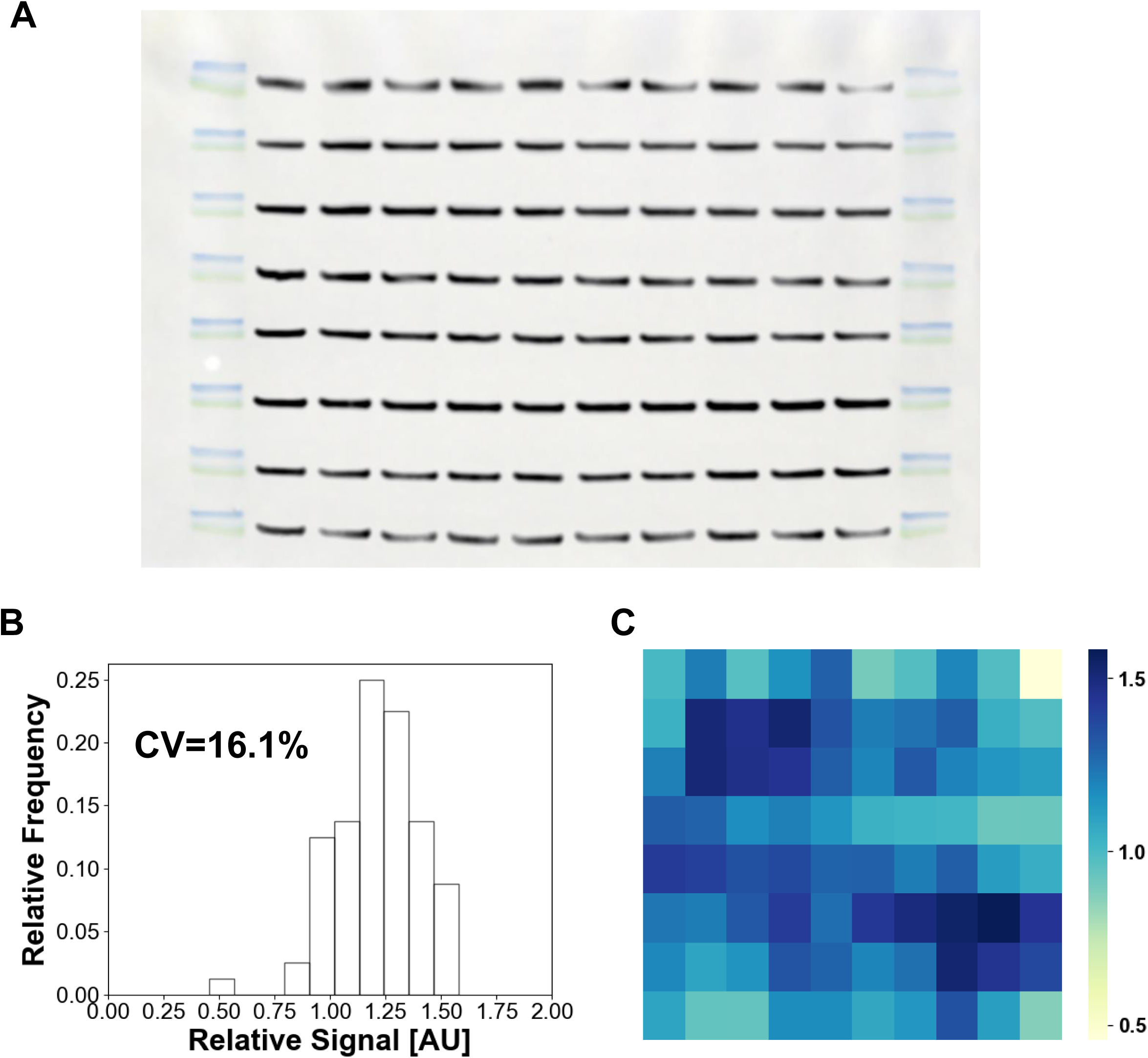
96-Well Western Blot with HEK293 Lysate and Analyzed for α-Tubulin. **A.** Full membrane scan with colorimetric ladder image embedded. A 96-well 10% Tris-HCl gel was loaded with 5 *μ*L of molecular weight ladder in each well of the first and last columns, and 20 *μ*L HEK293 lysate (see Methods) in the other wells. We then performed electrophoresis, transfer (30V, 16 hr, 4°C, nitrocellulose), and antibody incubations, and finally imaged. **B.** The bands from A were subjected to densitometry and the relative intensities analyzed for variation as a histogram. **C**. Densitometry results of band quantification from image in A. Each band corresponds to a square in the heatmap.

**Figure S4.**
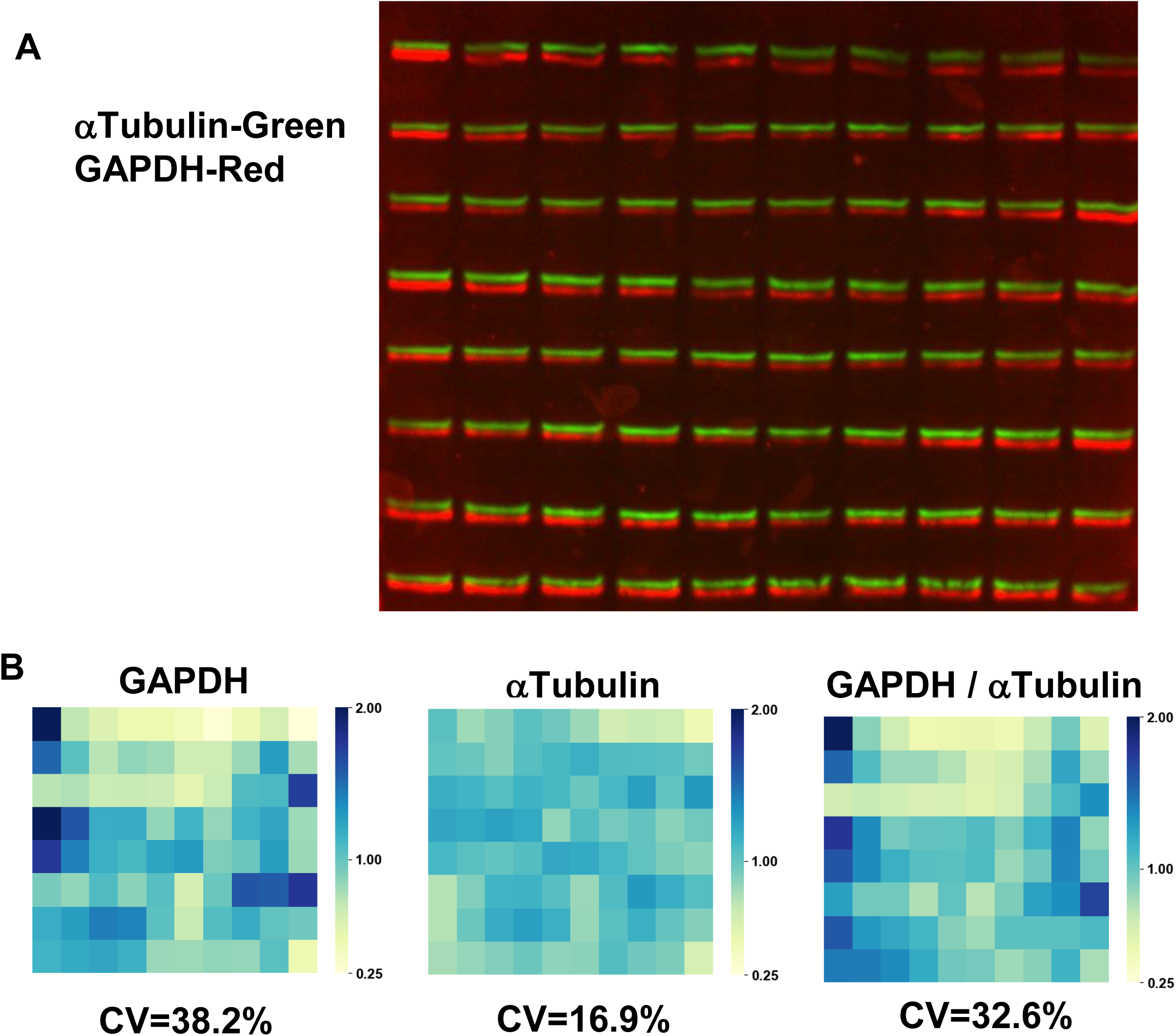
96-Well Western Blot with HEK293 Lysate and Analyzed for α-Tubulin and GADPH. **A.** Full membrane scan with Azure 600. A 96-well 10% Tris-HCl gel was loaded with 5 *μ*L of molecular weight ladder in each well of the first and last columns (colorimetric), and 20 *μ*L HEK293 lysate (see Methods) in the other wells. We then performed electrophoresis, transfer (30V, 16 hr, 4°C, nitrocellulose), and antibody incubations, and finally imaged. The ladder is not shown because it has extremely intense bands in the red pseudocolor channel. **B.** The bands from A were subjected to densitometry and the relative intensities analyzed. Each band corresponds to a square in the heatmap. The far right heatmp is the GAPDH intensities divided by the αTubulin intensities.

**Figure S5.**
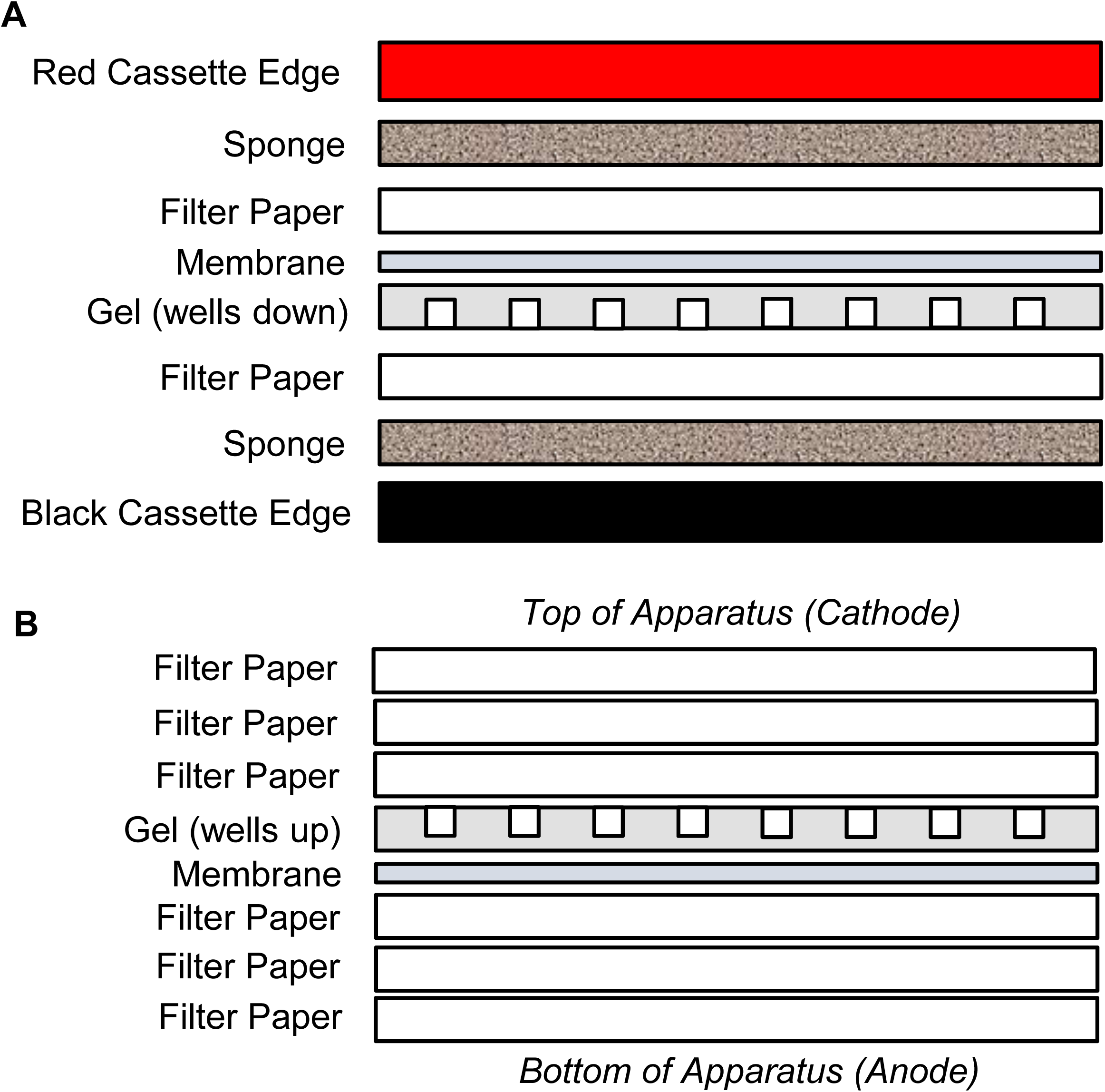
Transfer Sandwich Cartoons. **A.** Wet transfer sandwich. **B.** Semi-dry transfer sandwich.

